# GmPTF1 Modifies Root Architecture Responses to Phosphate Starvation in Soybean

**DOI:** 10.1101/830612

**Authors:** Zhaojun Yang, Ying He, Yanxing Liu, Yelin Lai, Jiakun Zheng, Xinxin Li, Hong Liao

## Abstract

Though root architecture modifications may be critically important for improving phosphorus (P) efficiency in crops, the regulatory mechanisms triggering these changes remain unclear. In this study, we demonstrate that genotypic variation in *GmEXPB2* expression is strongly correlated with root elongation and P acquisition efficiency, and enhancing its transcription significantly improves soybean yield in the field. Promoter deletion analysis was performed using six 5’ truncation fragments (P1-P6) of *GmEXPB2* fused with the *GUS* reporter gene in transgenic hairy roots, which revealed that the P1 segment containing 3 E-box elements significantly enhances induction of gene expression in response to phosphate (Pi) starvation. Further experimentation demonstrated that GmPTF1, a bHLH transcription factor, is the regulatory factor responsible for the induction of *GmEXPB2* expression in response to Pi starvation. In short, Pi starvation induced expression of *GmPTF1*, with the GmPTF1 product not only directly binding the E-box motif in the P1 region of the *GmEXPB2* promoter, but also activating *GUS* expression in a dosage dependent manner. Further work with soybean transgenic composite plants showed that, altering *GmPTF1* expression significantly impacted *GmEXPB2* transcription, and thereby affected root growth, biomass and P uptake. Taken together, this work identifies a novel regulatory factor, GmPTF1, involved in changing soybean root architecture through regulation the expression of *GmEXPB2*. These findings contribute to understanding the molecular basis of root architecture modifications in response to P deficiency, and, in the process, suggest candidate genes and a promoter region to target for improving soybean yield through molecular breeding of P efficiency.

**One Sentence Summary:** The bHLH transcription factor GmPTF1 regulates the expression of β-expansin gene *GmEXPB2* to modify root architecture, and thus promote phosphate acquisition, and biomass in soybean.

## INTRODUCTION

Phosphorus (P) is an essential mineral nutrient for plant growth and development. As a key structural component of biomolecules such as nucleic acids, proteins, and phospholipids, P is involved in multiple metabolic and biosynthetic processes required for the functioning of plant cells. Although the total amount of P in a given soil may be high, phosphate (Pi), which is the preferred form for assimilation, typically moves slowly through diffusion in the soil solution after being released from largely unavailable forms fixed to soil particles in aluminum-P, iron-P, and calcium-P bonds (Kochian et al., 2004; Rausch and Bucher, 2002). Low P availability significantly limits crop yields, and thus stands as a worldwide constraint for crop growth and productivity. Insights gained from better understanding the genetic and molecular mechanisms underlying plant adaptions to P deficiency, therefore, promise to spur development of smart crop cultivars that produce well in a range of P availability conditions through optimization of P utilization efficiency (Tian et al., 2012).

Plants have evolved a variety of complex responses and adaptations to P deficiency (Muneer and Jeong, 2015; Panigrahy et al., 2009). Notable examples include increasing accumulation of starch and anthocyanin (Chen et al., 2018; Leong et al., 2018; Wang et al., 2015), changing of root morphology and architecture (Gutierrez-Alanis et al., 2018; Li et al., 2016; Suen et al., 2018), enhancing Pi transport activity (Gu et al., 2016; Wang et al., 2017), and inducing endogenous and secreted phosphatases and RNases (Liang et al., 2002; Peng et al., 2018; Tian et al., 2014; Wang et al., 2009).

As one of the least available macronutrients, with very low mobility and high fixation in soils (Clarkson, 1981), Pi acquisition by plants often relies on the ability of root systems to most effectively explore the soil. The heterogenous distribution of Pi observed in many soils has led to discovery of plants with shallow root architectures from a soybean core collection, which provides an advantageous spatial frameworks for acquiring P from the P-rich topsoil (Zhao et al., 2004). Hence, root system architecture may be a critical component of efficient P acquisition in plants. Support for this model arises from multiple reports of Pi starvation stimulating the formation and emergence of lateral roots and root hairs (Bates and Lynch, 1996; Gaume et al., 2001; Williamson et al., 2001). In certain plant species, the formation of proteoid or cluster roots is another special type of root architectural adaptive response to P deficiency (Neumann and Martinoia, 2002; Zhou et al., 2008). Despite numerous reports of plants responding to Pi starvation with alterations in root morphology, the signaling and transcriptional regulation networks mediating these responses remain largely unknown.

Many genes involved in remodeling plant root system architecture in response to Pi starvation have been identified in a variety of plant species since the mid-aughts. For example, a diverse collection of transcription factors (TFs), including ZAT6, SIZ1, ARF7, ARF19, bHLH32 and WRKY75 from *Arabidopsis* (Chen et al., 2007; Devaiah et al., 2007a; Devaiah et al., 2007b; Huang et al., 2018; Miura et al., 2011), OsMYB5P and OsPHR2 from rice (Wu and Wang, 2008; Yang et al., 2018), and TaZAT8 from wheat (Ding et al., 2016), have been identified as playing critical roles in modifying root architecture in response to P deficiency. Interestingly, a large fraction of the identified Pi response regulators are members of the basic-helix-loop-helix (bHLH) family of transcription factors with their N-terminal basic region and a helix-loop-helix region (Li et al., 2006; Toledo-Ortiz et al., 2003). Several bHLH proteins have been shown to bind to E-box (CANNTG) *cis*-regulatory elements in the promoter regions of transcriptionally regulated downstream genes (Ito et al., 2012; Massari and Murre, 2000).

Several bHLH transcription factors associated with P deficiency responses have been discovered and characterized as regulators of root architecture remodeling in multiple crop species. In maize, the bHLH family member ZmPTF1 improves low Pi tolerance through regulation of carbon metabolism and root growth (Li et al., 2011). Recently, this gene, which binds to G-box (CACGTG) type E-box elements (Atchley et al., 1999; Massari and Murre, 2000), was associated with drought tolerance and found within the promoter regions of multiple drought stress responsive genes (Li et al., 2019). In rice, overexpression of *OsPTF1*, a bHLH transcription factor, significantly promoted increases in total root length and root surface area, which resulted in enhancement of Pi acquisition in comparisons with wild-type counterparts. Microarray analysis has further revealed a large set of Pi-starvation responsive genes that are up-/down-regulated by *OsPTF1* expression, and thus improve tolerance to Pi deprivation in rice (Yi et al., 2005). All of these findings suggest that bHLH members function as critical regulatory components in mediating root system architecture adaptations associated with P efficiency.

Soybean (*Glycine max*) is one of the most widely grown leguminous crops worldwide. Production, however, is often limited by soil P availability. Previously, we cloned and characterized a soybean β-expansin, *GmEXPB2*, from a Pi starvation induced cDNA library constructed from a P-efficient soybean genotype (Guo et al., 2008; 2011). *GmEXPB2* appears to be primarily expressed in roots and dramatically induced by Pi starvation. Overexpression of *GmEXPB2* significantly promotes root elongation, and is accompanied by increases in plant growth and P uptake under P deficiency growth conditions (Guo et al., 2011; Zhou et al., 2014). Genetic modification of root morphology and architecture might, therefore, be effective strategies for improving crop production in Pi limited soils. However, transcriptional regulators of *GmEXPB2* responses to Pi starvation remain unknown.

In the present study, we take *GmEXPB2* as our Pi deficiency response subject. This β-expansin gene is known to be critically involved in root system architecture responses to Pi starvation in soybean (Guo et al., 2011). Since *GmEXPB2* is primarily expressed in roots and dramatically induced by Pi starvation, its expression is likely controlled by a transcriptional factor possibly binding to P responsiveness *cis*-elements. However, none of the motifs associated with P efficiency to date have yet been identified and functionally analyzed in the promoter region of *GmEXPB2*. To understand the molecular basis of the low P stress response and identify ideal candidate promoters for transgenic breeding of P efficiency, the *GmEXPB2* promoter region was analyzed by testing a set of mutants harboring a systematic series of deletions for responses to low P availability. Then, the *cis*-elements identified as regulated by the transcription factor GmPTF1 were assayed in transgenic tobacco leaves and hairy roots. Participation of GmPTF1 in root growth and its contributions to P efficiency was further confirmed in soybean transgenic composite plants, as was induction of *GmEXPB2* transcription. The results presented here confirm that *GmEXPB2* acts in root growth and yield responses to Pi starvation, and further demonstrates that this vital component of P efficiency in soybean is activated by the bHLH type transcription factor GmPTF1.

## RESULTS

### Genotypic Variation in *GmEXPB2* Expression and Association with Root Elongation and P Acquisition Efficiency

To investigate whether the genotypic variation observed for *GmEXPB2* expression is associated with root elongation and P efficiency, 111 genotypes from a soybean core collection (Zhao et al., 2004) were classified into three groups according to the relative expression level of *GmEXPB2* in roots under -P conditions. These were labeled group I, II and II with low, intermediate and high expression levels of *GmEXPB2*, respectively. We discovered that the root length and P acquisition efficiency varied among these groups, with group III genotypes exhibiting the highest expression of *GmEXPB2*, as well as, the longest roots and highest P contents of any of the three groups (Fig. 1). This result suggests that enhancement of *GmEXPB2* expression might contribute to root growth and improvement of P efficiency.

**Figure 1.**
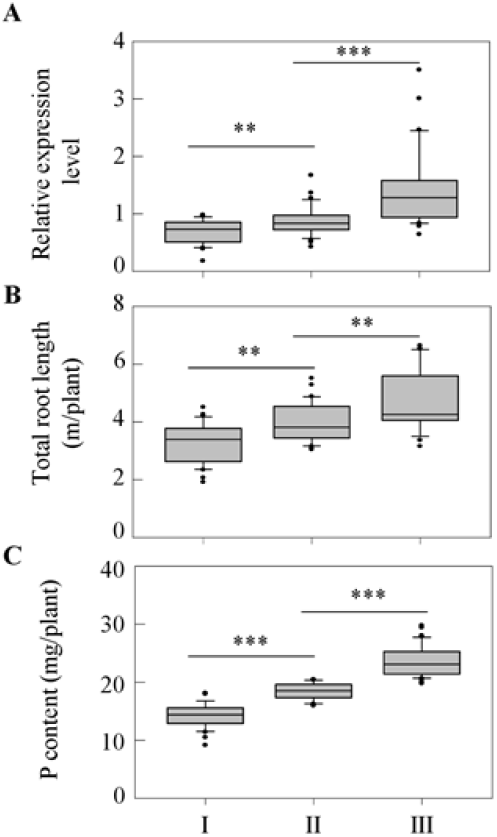
Association of *GmEXPB2* expression levels observed across a soybean core collection with P acquisition efficiency and root growth, The 111 observed soybean genotypes were categorized into three groups according to *GmEXPB2* transcription in roots under low P conditions. I, II, and III represent lower, intermediate, and higher expression level categories of *GmEXPB2*, respectively. P acquisition efficiency was calculated as total P content per plant. Asterisks represent significant differences between groups for the same tissue in the Studen’s *t* test (**: 0.001 < P ≤0.01, ***: *P* ≤0.001).

### Overexpression of *GmEXPB2* Significantly Improves Soybean Yield through Promotion of Root Growth and P Uptake

We further performed field trials to evaluate the effects of *GmEXPB2* expression on soybean yield. In these experiments, overexpression of *GmEXPB2* (OE) significantly improved soybean growth and yield (Fig. 2A). In comparisons with wild type (WT) plants, three OE lines produced 12.1, 23.8 and 30.4% increases in pod number (Supplemental Fig. S1A), 12.1, 20.7 and 24.4% increases in seed number (Fig. 2B), 18.1, 25.5 and 27.6% increases in grain weight (Fig. 2C), and 10.0, 7.9 and 5.2% increases in 100 grain weight (Supplemental Fig. S1B).

**Figure 2.**
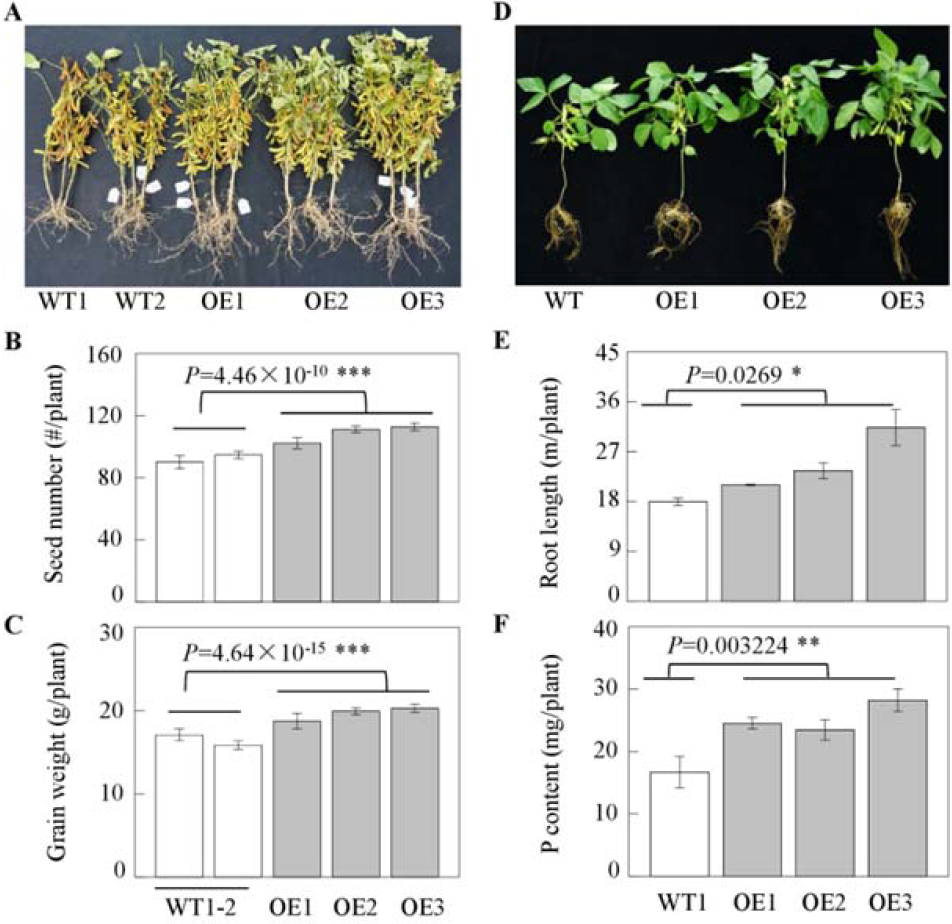
Overexpressing *GmEXPB2* significantly improved soybean root growth and P acquisition efficiency in a soil pot experiment, as well as, yield in a field trial, A, Growth performance B, Total root length, C. Plant P content. D, Photograph showing soybean growth performance in a field trial. E, Seed number. F, Grain weight. Total root length and plant P content were measured at the R5 stage WT: wild-type plants; OE: *GmEXPB2* overexpression lines. Data are means of 4 biological replicates with SE in the pot experiment, and means of 40-90 independent plants with SE in the field trial. Asterisks represent significant differences between groups in the Student’s *t* test (*: 0.01 < *P* ≤ 0.05,**: 0.001 < P ≤ 0.01, ***: *P* ≤ 0.001).

Further investigation of how *GmEXPB2* expression affects root growth and P efficiency was conducted with three OE lines of *GmEXPB2* and WT plants grown in pots to the R5 stage (Fig. 2D). After confirming the presence of the *bar* gene through qualitative PCR, accumulation of *GmEXPB2* mRNA was monitored in leaves RT-qPCR. In these tests, expression of *GmEXPB2* was 3.54, 5.71, and 2.33-fold higher in the three OE transgenic lines than in WT plants (Supplemental Fig. S2). Overexpression of *GmEXPB2* significantly promoted soybean root elongation and thus P acquisition efficiency. Relative to WT plants, the overexpression of *GmEXPB2* led to 17.2%, 31.4%, and 74.9% increases in total root length, along with 46.9%, 40.5%, and 69.0% increases in P content (Fig. 2, E and F). These results suggest that increasing *GmEXPB2* expression improves P efficiency through regulation of adaptive changes in root system architecture, which ultimately leads to increases in soybean yield in field.

### Phosphorus Availability Regulates *GmEXPB2* Expression in Different Tissues

Previous studies have demonstrated that *GmEXPB2* expression is not only involved in root system architecture responses to Pi starvation (Guo et al., 2011), but also improves nodulation regardless of P availability (Li et al., 2015). To characterize the temporal and spatial patterns of *GmEXPB2* expression in response to Pi starvation, roots, nodules and leaves were collected from plants grown in P deficient or sufficient conditions and separately assayed for *GmEXPB2* transcription. The results clearly show that *GmEXPB2* transcripts were predominantly localized to roots and were up-regulated by P deficiency (Fig. 3A). At 7 days after inoculation (DAI), *GmEXPB2* was most abundantly expressed in nodules, followed by bulk roots, but not in leaves. Then, by 14 DAI, *GmEXPB2* expression was significantly enhanced in roots, especially under Pi deficiency conditions, as well as, in nodules. These results confirm that *GmEXPB2* is indeed involved in nodule development during early stages of organogenesis (Li et al., 2015). Plus, we reach the additional conclusion here that *GmEXPB2* also appears to primarily participate in adaptive changes of roots responding to P deficiency, especially under long term Pi deficiency conditions.

Further tissue localization of *GmEXPB2* transcripts in soybeans subjected to P deprivation was conducted using GUS staining of soybean transgenic composite plants carrying the promoter region of *GmEXPB2* fused to the β-glucuronidase (GUS) reporter gene *(proGmEXPB2*::GUS) grown under Pi starvation conditions (Fig. 3B). Consistently, *GmEXPB2* reporter expression was significantly induced in roots by P deficiency conditions at 14 DAI relative to 7 DAI. Meanwhile, in nodules, GUS staining from the *proGmEXPB2*::GUS construct was strongest during early development at both 7 and 14 DAI. Taken together, these results indicate that *GmEXPB2* might play distinct roles in different tissues, with regulation of root growth responses to limited Pi availability predominating over participation in nodule development.

**Figure 3.**
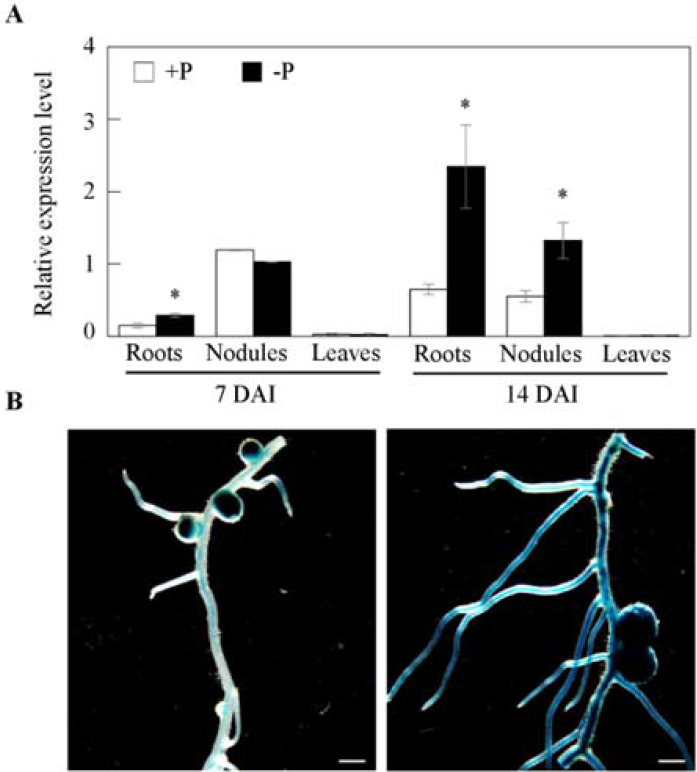
Regulation of *GmEXPB2* expression by P supply. A, The expression of *GmEXPB2* in roois, leaves, and nodules was determined at 7 and 14 days after inoculation (DAI) with rhizobia under -P and +P conditions. The -P and +P treatments included 5 and 200 μM KH_2_PO_4_, respectively. Data are means of 3 replicates with SE. Asterisks represent significant differences in *GmEXPB2* expression in a given tissue between -P and +P treated plants as determined in the Student’s *t* test {*: 0.01 < *P* ≤ 0.05). B, GUS staining in transgenic hairy roots and nodules developing in -P nutrient solution. Soybean transgenic composite plants harboring *proGmEXPB2*:: GUS were grown in sand culture irrigated with -N and -P nutrient solution for 7 and 14 DAI with гhizobia. Scale bar=2 mm.

### Identification of P Responsive Regulatory Segments of the *GmEXPB2* Promoter Region in Transgenic Soybean Composite Plants

To characterize functional components of the *GmEXPB2* promoter, six deletion fragments were separately fused to the *GUS* reporter gene and transferred into hairy root transformants by hypocotyl injection. Promoter deletions were designated as P1 to P6, with each including different lengths of promoter sequence from the translation start codon (ATG) of *GmEXPB2* (Fig. 4A). All of the tested truncated promoters were able to drive *GUS* gene expression in transgenic hairy roots (Fig. 4B). However, only the roots harboring P1- and P2-promoter segments exhibited obvious induction of *GUS* expression in response to Pi starvation. Specifically, in response to P deprivation, the respective increases in *GUS* expression and its protein activity were 3.25- and 2.83-fold for P1 plants, and 4.67- and 1.71-fold for P2 plants (Fig. 4, C and D). No other deletion lines exhibited distinct differences in GUS expression/activity between -P and +P conditions. These results suggest that the P1 promoter (−304 to −1 bp) region contains Pi starvation responsive elements that induce *GmEXPB2* expression, while as yet unidentified repressors act on elements around 500 bp upstream or more from the start codon.

**Figure 4.**
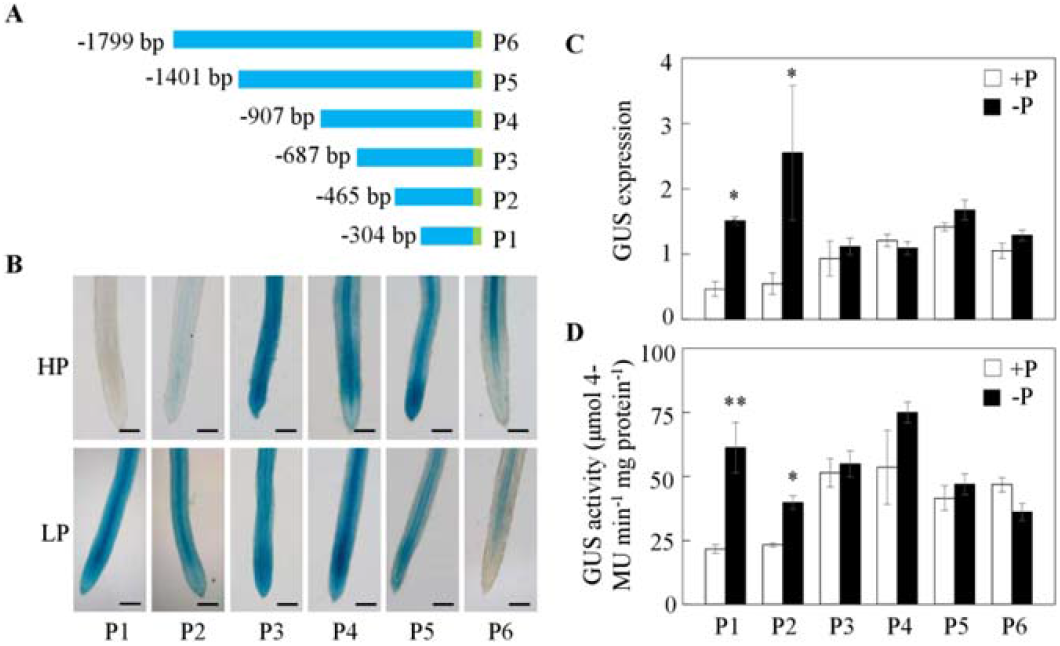
Deletion analysis of the *GmEXPB2* promoter. A, Schematic outlines of the truncated *GmEXPB2* promoters (P1 to P6) fused with the *GUS* reporter gene. B, GUS staining of hairy roots transformed with the indicated constructs under -P and +P conditions. Scale bar=l mm. C, Relative expression of the *GUS* gene D, Quantitative GUS activity analysis of the transgenic hairy roots by fluorimetric assay Data are means of 5 replicates with SE. Asterisks represent significant differences between promoters from PI and P2 for the same trait in Student’s *t* tests {*: 0 01 < *P* ≤ 0 05, **: 0.001 < *P* ≤0.01).

### *Cis*-regulatory Elements Identified in the *GmEXPB2* Promoter

In order to characterize the Pi starvation responsive *cis*-regulatory elements of the P1 promoter, putative *cis*-elements were first identified in a search of the NEW PLACE database (https://www.dna.affrc.go.jp/PLACE/?action=newplace). This returned a total of 40 bonding sites distributed unequally throughout the 304 bp upstream region of *GmEXPB2* (Supplementary table S3). Among these identified putative *cis*-elements, four are known for tissue-specific gene expression including ROOTMOTIFTAPOX1 (AATAT), RHERPATEXPA7 (ACGTGA), NODCON1GM/OSE1ROOTNODULE (ATCTTT) and POLLEN1LELAT52 (AGAAA), with root, root hair, nodulin and pollen specific motifs, respectively (Elmayan and Tepfer, 1995; Filichkin *et al*., 2004; Kim *et al*., 2006; Stougaard *et al*., 1990). Several environmental stress-related motifs were also found in the P1 promoter. For example, ACGTATERD1 is required for etiolation-induced expression (Simpson *et al*., 2003), while GATABOX and GT1CONSENSUS are mainly light-responsive regulatory elements (Rubio-Somoza *et al*., 2006; Zhou, 1999), and MYBCORE is chiefly responsive to water stress (Urao *et al*., 1993). In addition, three EBOXBNNAPA/MYCCONSENSUSAT/MYCATRD22 (E-box, CANNTG) motifs were identified at positions 208, 215 and 251 of the *GmEXPB2* upstream sequence (Supplementary table S3; Fig. 5A). The E-box sequence can bind basic-helix-loop-helix (bHLH) type transcription factors, and GmPTF1, a bHLH family member, is a known mediator of tolerance to Pi deprivation in soybean (Li *et al*., 2014; Massari and Murre, 2000). In this context, GmPTF1 can be considered as a candidate regulating factor of soybean root architecture modification responses to Pi starvation, with responses arising primarily via transcriptional regulation of *GmEXPB2*.

**Figure 5.**
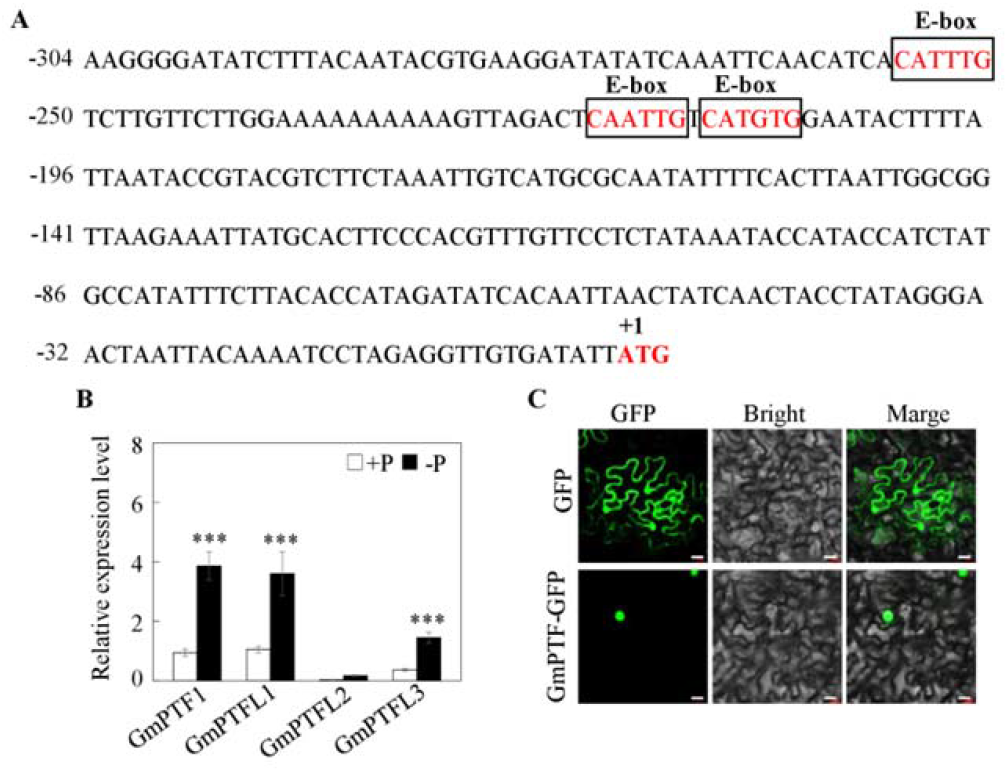
E-box elements in the promoter of *GmEXPB2* and putative regulatory transcription factors in soybean. A, Relative positions of the E-boxes. The translational start codon ATG was assigned position +1, and the numbers flanking the sequences of the *GmEXPB2* promoter fragments were counted from there The E-boxes are indicated within black rectangles. B, Transcripts of *GmPTFs* in soybean roots growing under -P and +P conditions Data are means of 5 replicates with SE. Asterisks represent significant differences between -P and +P for the same gene in the Student’s *t* test (***: *P* ≤0.001). C, Subcellular localization of GmPTF1 fused to GFP in tobacco cells. Cells were observed by green GFP fluorescence of the GmPTF1 protein (GmPTF1-GFP) with a GFP empty vector driven by 35S included as control vectors. Scale bar=100 μm.

### Characterization of *GmPTF* Genes and Subcellular Localization of GmPTF1

Although GmPTF1 is known as a homologue of OsPTF1 that was isolated from soybean years ago (Li et al., 2014), any functionality of this gene in soybean responses to Pi starvation remains largely unclear. Here, we first quantified the extent of the GmPTF family in the soybean genome through a search of the phytozome website (http://www.phytozome.net/). This returned 170 *GmbHLH* genes in the soybean genome. They are unevenly distributed on all chromosomes from 1 to 20. A phylogenetic tree was further constructed by neighbor-joining analysis in MEGA 4.1 to determine the evolutionary relationships among the GmbHLHs family members (Supplementary Fig. S3). This demonstrated that soybean GmbHLH proteins sort into six distinct groups. Most of the previously tested GmbHLHs belong to group III, which includes 42 of the 170 bHLH members. GmPTF1 (Glyma19G143900) and its three homologs are also group III bHLHs.

To elucidate how this subset of GmPTF1 homologs respond to P deficiency, RT-qPCR analysis was carried out using total RNA from 25 d-old roots (Fig. 5B). Other than *GmPTFL2*, which was barely detected in either P treatment, the remaining GmPTF1 homologs, *GmPTF1, GmPTFL1* and *GmPTFL3*, were significantly up-regulated by more than 4.1-, 3.4- and 3.9-fold, respectively, in response to Pi starvation. Given that *GmPTF1* exhibited the strongest response to Pi deprivation, and that previous research suggests that *GmPTF1* contributes to tolerance of Pi starvation in soybean (Li *et al*., 2014), *GmPTF1* was selected as candidate gene for further study.

To define the subcellular localization of GmPTF1, tobacco leaves were infiltrated with *agrobacterium tumefaciens* harboring a GmPTF1-GFP fusion. In contrast to control expression of GFP alone, which distribute throughout the nucleus and cytosol, the GFP signal derived from the fusion was exclusively confined to the nucleus (Fig. 5C). This is consistent GmPTF1 acting as a transcription factor in nucleus where it might regulate downstream gene transcription.

### Dosage Effects of E-box Elements in the *GmEXPB2* Promoter

Since GUS activity was significantly enhanced by Pi starvation of plants with the P1 promoter containing three E-box elements, and with *GmPTF1* known as an E-box binding transcription factor, the effects of *GmPTF1* expression on *GmEXPB2* promoter driven GUS activity were investigated in an *Agrobacterium*-mediated co-transient assay performed in tobacco leaves (Fig. 6). The P1 promoter was further truncated and named pro1 (−213 to −1) and pro2 (−220 to −1), harboring one and two E-boxes, respectively (Fig. 6A). Although *GUS* was expressed with all three promoters, *GmPTF1* only up-regulated *GUS* in co-transgenic tobacco leaves also harboring pro2::GUS or P1::GUS (Fig. 6B). Transcription of the *GUS* gene and GUS activity itself exhibited respective increases of 49% and 133% for P1, and 48% and 102% for pro2 when co-transformed with *GmPTF1* in tobacco leaves (Fig. 6, C and D). In short, there appears to be a dosage effect of E-boxes on the transcription of *GmEXPB2* regulated by GmPTF1. At least two E-boxes are necessary for *GmEXPB2* expression to be altered by GmPTF1. Three E-boxes may promote more responsive *GmEXPB2* expression than two.

**Figure 6.**
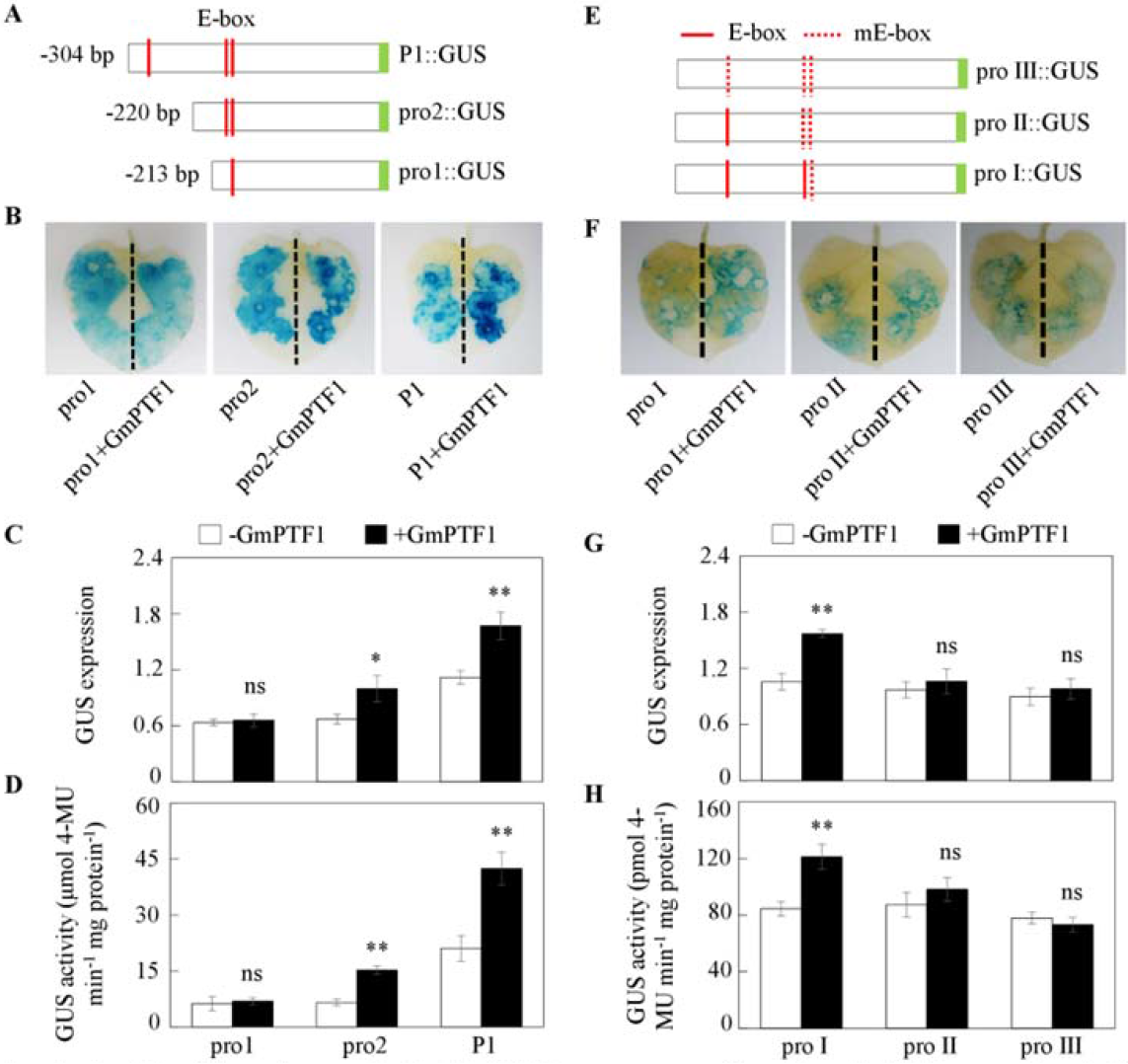
Functional analysis of the E-box elements in the *GmEXPB2* promoter as affected by *GmPTF1* expression A, Schematic outlines showing the *GmEXPB2* promoter harboring different numbers of normal (E) or mutated E-boxes (niE-box). B and F, GUS staining of tobacco leaves. C and G, Relative expression of the *GUS* gene. D and H, Quantitative GUS activity analysis of transgenic tobacco leaves in fluαrimetric assays. Data are means of 6 replicates with SE. Asterisks represent significant differences between promoters with different numbers of E-boxes for the same trait in the Student’s *t* test (*: 0.01 < *P* ≤ 0.05, **: 0.001 < *P* ≤ 0.01). ns, Not significant at the P=0.05 threshold.

To further confirm the effects of *GmPTF1* on P1 promoter mediated GUS activity, three expression vectors containing E-boxes modified from the *GmEXPB2* promoter were constructed and investigated (Fig. 6E). Interestingly, GUS staining was obviously detectable in all tobacco leaves containing *GUS* fused to a *proI, proII*, or *proII* promoter (Fig. 6F). However, GUS activity was induced by *GmPTF1* expression only in leaves co-transformed with proI::GUS.

Co-expression of *proI* and *GmPTF1* led to 48.7% and 43.4% increases in *GUS* expression and its protein activity, respectively (Fig. 6, G and H). Together, these results suggest that the E-box element of the *GmEXPB2* promoter is required for *GmPTF1* activation, and increasing the number of E-box elements in the *GmEXPB2* promoter may boost the expression of the regulated gene.

### Alteration of *GmPTF1* Expression Influences Transcription of *GmEXPB2*, and Thereby Promotes Root Architecture Modifications in Transgenic Soybean Composite Plants

In order to better understand whether *GmPTF1* expression might affect the transcription of *GmEXPB2*, and thus root growth in soybean plants, *GmPTF1* overexpression and RNA interference lines (OE and Ri lines) of transgenic composite plants were generated. The quality of gene transformation in transgenic hairy roots was checked through GFP fluorescence microscopy and RT-qPCR analysis. After selecting transgenic hairy roots under a microscope (Supplementary Fig. S4), one transgenic hairy root per plant was used for further study. In RT-qPCR analysis, transcription of *GmPTF1* was 49.8 times higher in OE and 1.87 times lower in Ri plants than that in CK lines (Fig. 7A). Moreover, altering *GmPTF1* expression significantly influenced the expression of *GmEXPB2* in roots, which increased by 40.1% in OE plants, and decreased by 60.5% in Ri plants when compared with CK lines (Fig. 7B). Furthermore, *GmPTF1* expression significantly affected soybean root growth in transgenic composite plants, with *GmPTF1* overexpressing hairy roots growing much better than control lines expressing the empty vector, and suppression of *GmPTF1* producing the opposite effects (Fig. 7C). After 25 days, roots of *GmPTF1* overexpressing transgenic composite plants were 47.1% longer, and those of Ri lines were 36.4% shorter than control roots (Fig. 7D). These findings indicate that *GmPTF1* regulates soybean hairy root growth through effects on the expression of *GmEXPB2*.

**Figure 7.**
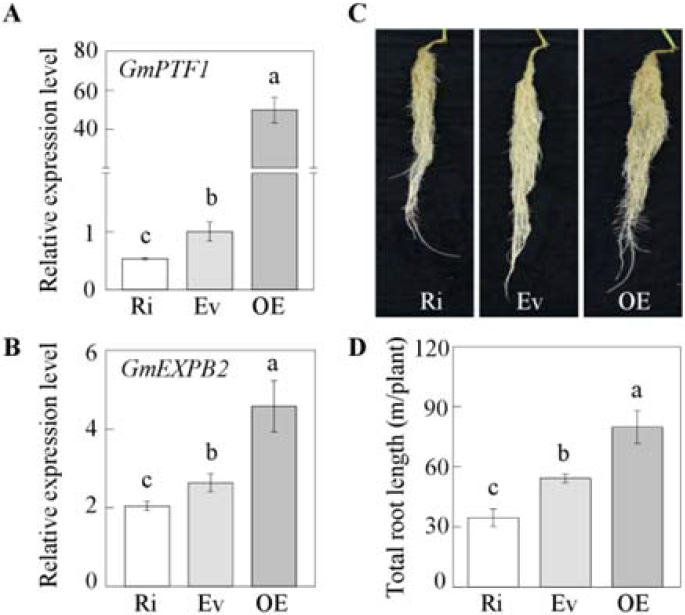
*GmPTF1* regulation of root growth acting through effects on *GmEXPB2* expression in transgenic composite soybean plants. A, *GmPTF1* expression in hairy roots B, *GmEXPB2* expression in *GmPTF1* overexpressing (OE) or RNA interference (Ri) transgenic lines. C, Growth performance of hairy roots, D, Total root length. Composite soybean plants with transgenic hairy roots were grown in nutrient solution for 25 d before separately harvesting roots for analysis. Ev: control transgenic soybean hairy roots harboring empty vector Each soybean transgenic composite plant represents one independent transgenic line, and one independent transgenic plant was considered as one biological replicate. Relative expression was normalized against the geometric mean of Ev transcription. Data are means of 5 replicates with SE. Different letters indicate significant differences between Ri or OE lines and Ev control plants for the same trait in a two-way ANOVA test (*P* < 0.05).

### *GmPTF1* Expression Enhances Plant Growth and P Content in Soybean Transgenic Composite Plants

Impacts of *GmPTF1* on plant growth and P efficiency were further investigated in soybean transgenic composite plants. Soybean growth was enhanced in OE lines and inhibited in Ri lines in comparisons with CK plants (Fig. 8A). Overexpression of *GmPTF1* led to increases of 57.9%, 46.2% and 30.7% in root dry weigh, plant fresh weigh and P content, respectively. Meanwhile, suppression of *GmPTF1* resulted in a 50.5% decline in root dry weight, along with 34.6% and 49.5% declines in plant fresh weight and P content, respectively (Supplementary Fig. S5; Fig. 8, B and C). These results indicate that regulatory effects of *GmPTF1* ultimately play important roles in overall root growth and efficiency of P utilization.

**Figure 8.**
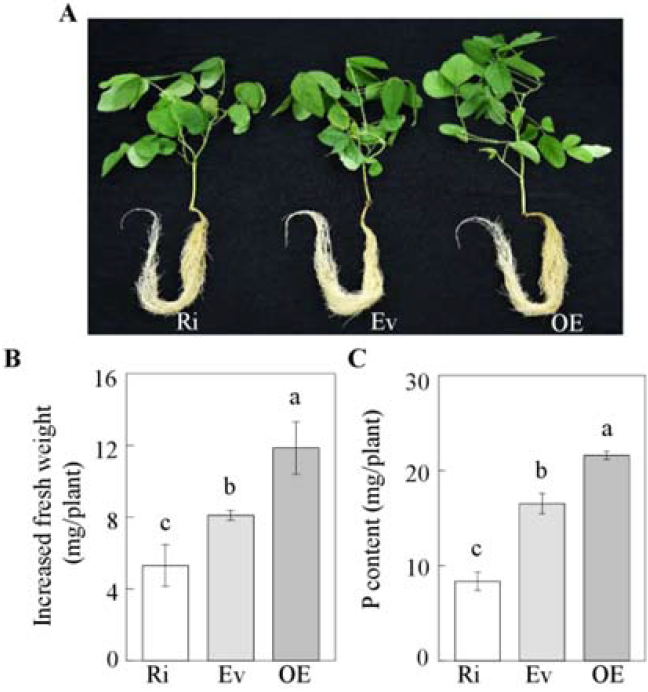
Effects of *GmPTF* expression on growth of soybean transgenic composite plants. A, Phenotype of composite soybean plants. B. Plant fresh weight, C, P content, Composite soybean plants with transgenic hairy roots were grown in normal nutrient solution for 25 d. Ev: control transgenic soybean noduies harboring empty vector; OE: *GmPTF* over-expressing transgenic lines; Ri: *GmPTF* RNA interference transgenic lines. Each soybean transgenic composite plant represents one independent transgenic line, and one independent transgenic plant was considered as one biological replicate, Data are means of 5 replicates with SE, Different letters indicate significant differences between Ri or OE lines and Ev control plants for the same trait in a two-way ANOVA test (*P* < 0.05).

## DISCUSSION

Low phosphorus (P) availability is a major constraint on plant growth and production worldwide. As the main organ involved in the acquisition of nutrients and water, roots are a logical subject of research efforts aiming to incorporate plant adaptations for growth in P limited soils into crops efficiently acquiring and utilizing P while maintaining high yields. Although a series of genes have been reported as involved in root architecture responses to Pi starvation, to date, the transcriptional regulatory mechanisms underlying these responses has remained largely unknown. Overexpression of *GmEXPB2* is known to significantly enhance root growth and Pi uptake (Guo et al., 2011). Here, we further evaluated how *GmEXPB2* expression influences P efficiency in soybeans selected from a diverse core collection (Fig. 1). Interestingly, across the genotypes tested, variation in *GmEXPB2* expression was strongly correlated with root elongation, P acquisition efficiency, and resulting soybean yields in field studies (Figs. 1 and 2; Supplementary Fig. S1). This indicates that *GmEXPB2* is indeed an important contributor to P efficiency and maintenance of yield through modifications in root system architecture. It also suggested that *GmEXPB2* would be an interesting subject in detailed exploration of transcriptional regulatory pathways guiding P deprivation responses in soybean.

In addition to impacting P efficiency, overexpression of *GmEXPB2* also improved soybean nitrogen efficiency through facilitation of nodulation, which also led to modifications in root architecture regardless of P supply (Li et al., 2015). Given that *GmEXPB2* transcripts were most abundant in early stages of nodule development, this gene was also tested for transcriptional responses to P deficiency in roots, nodules, and leaves at 7 and 14 DAI in RT-qPCR analysis and GUS staining assays (Fig. 3). Consistently, *GmEXPB2* was found to be predominantly expressed in young nodules under both -P and +P conditions, but was also highly induced by Pi starvation in roots. This result was also supported by the observation of GUS staining in soybean transgenic composite plants carrying the *proGmEXPB2*::GUS (Fig. 3B). Further experiments in this study, therefore, focus on outlining the molecular mechanisms regulating the accumulation of *GmEXPB2* mRNA in roots responding to Pi starvation. Moreover, the expression of *GmEXPB2* in nodules at 14 DAI was significantly enhanced by P deprivation. This result stands in contrast to a previous report (Li et al., 2015), in which P levels did not affect *GmEXPB2* expression in nodules at either 7 or 14 DAI. It is possible that *GmEXPB2* expression in nodules varies with differences in growth conditions or developmental stages between the previous report and this work.

A number of studies have demonstrated that gene promoters are important mediators of gene expression responses to stress and developmental processes (Sharma et al., 2017; Timerbaev and Dolgov, 2019; Zhang et al., 2017). To investigate how *GmEXPB2* transcription in roots was induced by P deprivation, a set of promoter deletions was analyzed in transgenic composite hairy roots. Among tested promoter segments, the P1 and P2 sequences (304 and 465 bp upstream sequences from translation start codon ATG) contained the key region for induction of *GmEXPB2* expression in response to Pi starvation (Fig. 4). This result confirms histochemical expression patterns reported previously in transgenic hairy roots carrying a 500 bp promoter sequence of *GmEXPB2* (Guo et al., 2011), which suggests that this fragment harbors P deficiency response activators. More precisely, the P1 associated strong responses in GUS activity to P deprivation indicate that the main *cis*-regulatory elements responsible for P deficiency responses lie between positions −304 and −1 bp.

Interestingly, no significant effects of P were observed in the roots of P3-P6 (587-1799 bp) plants (Fig. 4). An as yet unknown low P response inhibitor might be located within the 587 to 1799 bp sequence shared by P3-P6. This would allow for transcription of *GmEXPB2* to be influenced by multiple competing stimuli and fine tuned for a range of conditions. This is in accordance with the model of a promoter region as a collection of diverse transcription factor binding sites coordinating specific responses to complex sets of stimuli that may include hormonal, physiological, or environmental cues (Wray *et al*., 2003). Given the potential complexity of signals impacting the *GmEXPB2* promoter, a comprehensive description of the mechanisms guiding *GmEXPB2* responses to Pi starvation at the transcriptional level requires further study.

Many studies have also documented the involvement of *cis*-regulatory elements in a variety of regulatory networks adapted to mediate responses to fluctuating physiological and environmental conditions (Hanifiah et al., 2018; Kim et al., 2006; Rawat et al., 2005). In one relevant example, the R2R3 MYB transcription factor PHR1 and its homologs appear to play central roles in P signaling and Pi homeostasis (Bari et al., 2006; Chiou and Lin, 2011; Liang et al., 2013; Sun et al., 2016; Xue et al., 2017). PHR1 binds to the P1BS element (GNATATNC) in the promoter region of multiple Pi starvation induced genes, which activates gene expression (Franco-Zorrilla et al., 2004; Rubio et al., 2001). Site-specific mutation or deletion of the P1BS element in the promoter region abolished the transcription of a number of P responsive genes (Karthikeyan *et al*., 2009; Oropeza-Aburto *et al*., 2012).

In the current study, no P1BS elements were found in the 304 bp or 2 kb promoter fragments of *GmEXPB2*, while the three E-box elements described herein were closely associated at positions 208, 215, and 251 bp upstream of the *GmEXPB2* start codon (Supplementary Table S3; Fig. 5A). bHLH transcription factors are known to bind to E-box sequences (Ito et al., 2012; Massari and Murre, 2000). While a total of 170 putative bHLH members were identified in the soybean genome, *GmPTF1* displayed the largest increase of transcript levels in roots subjected to -P stress (Supplementary Fig. S3; Fig. 5B). Further GUS staining and quantitative analysis in transient co-transgenic tobacco leaves revealed that E-boxes were required for GmPTF1 to induce transcription of *GmEXPB2*, furthermore, this induction appears to depend on the number of E-boxes in a dosage-dependent manner (Fig. 6). This regulation of *GmEXPB2* by GmPTF1 was confirmed in observations of *GmEXPB2* expression being up- or down-regulated in soybean transgenic composite lines overexpressing or suppressing of *GmPTF1* (Fig. 7, A and B). Furthermore, altering *GmPTF1* expression significantly impacted root growth, plant biomass and P content (Figs. 7 and 8), which could be partially reproduced through manipulation of *GmEXPB2* transcription in hairy roots.

In summary, the results presented here demonstrate that GmPTF1 modifies root architecture responses to Pi starvation in soybean. Accordingly, we developed a schematic representation for how *GmEXPB2* is involved in root growth and yield as regulated by GmPTF1 (Fig. 9). As a bHLH transcription factor, GmPTF1, may recognize E-box binding sites in the promoter of *GmEXPB2*, which are necessary for P responsive changes in *GmEXPB2* transcription in roots. From there, downstream effects may include root elongation and root architecture modifications that allow for soybean plants to acquire more P and, ultimately, improve soybean biomass and yield. In general, the results described in this work will be useful for understanding the molecular regulation of genes involved in tolerance to Pi deprivation through effects on root architecture. With further study, this report might be useful for producing high yielding soybeans bred for enhanced efficiency of P acquisition and utilization.

**Figure 9.**
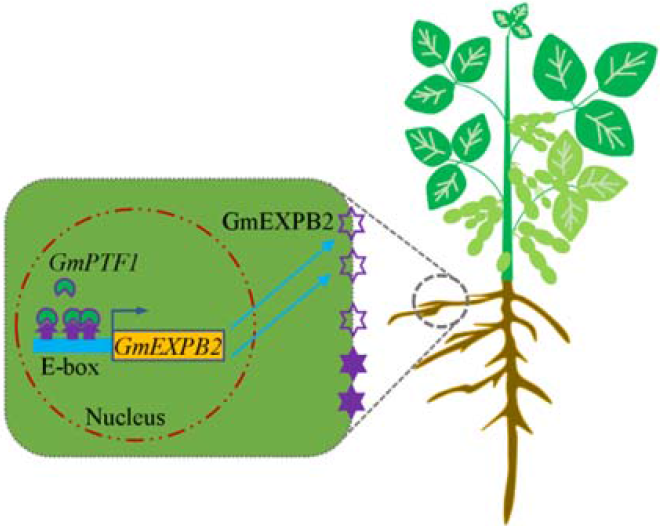
Hypothetical model of the *GmEXPB2* expression regulated by GmPTF1. In plants, the GmPTF1 directly enhanced *GmEXPB2* expression possibly by binding to the E-box motifs within the promoters of *GmEXPB2* to facilitate cell wall loosing, and thus modify root architecture and promote soybean yield.

## MATERIALS AND METHODS

### Plant Material and Growth Conditions

Experiments in this study included plants grown in the field, pots and hydroponics. For the field experiment, 111 genotypes from the soybean core collection were grown at the Boluo experimental farm (E114.28°, N23.18°) of South China Agricultural University, Huizhou City, Guangdong Province (Zhao et al., 2004). At the seed-filling stage, plants were harvested to analyze total root length and P content.

To study whether overexpression of *GmEXPB2* improves soybean yield, a field experiment was conducted in 2016 at the Ningxi experimental farm (23°130N, 113°810E) of South China Agricultural University, Guangzhou City, Guangdong Province. The basic soil chemical properties have been previously outlined (Wang et al., 2009). Seeds of wild type (WT) and three independent T_4_ stably transgenic lines overexpressing *GmEXPB2* (OE) were inoculated with rhizobia and then grown from March to June. There were three plots of each line, and 30 seedlings in each plot. Fifteen days after sowing, transgenic plants were identified in leaf painting herbicide assays. Seeds were harvested at the maturation stage for recording pod and seed number, along with seed weight after air-drying.

For the soil pot experiment, WT and three OE lines were germinated on vermiculite for 5 days prior to transplanting uniform seedlings into pots. The basic soil chemical characteristics were as follows: a pH of 6.46, 72.80 mg·kg^-1^ available N, 67.49 mg·kg^-1^ available P, and 2.7% organic matter. After 15 d of growth, leaves were harvested for *bar* gene identification and real-time quantitative reverse transcription PCR (RT-qPCR) analysis. At the R5 stage, plants were separately sampled for total root length and P content measurements.

For the hydroponic experiment, seeds from the soybean core collection were surface sterilized in 3% H_2_O_2_ for 1 min, rinsed with distilled water, and germinated in vermiculite for 7 d. Uniform seedlings were cultured in soybean growth solution with low Pi supplied as 5 μM KH_2_PO_4_ as describe previously (Qin et al., 2012). Plants were grown in growth chambers (day/night: 14 h/11 h, 26°C/24°C) for 14 d. Roots were harvested for RT-qPCR assays to test for a relationship between *GmEXPB2* expression and root elongation and P efficiency.

To study temporal and spatial patterns of *GmEXPB2* expression in response to Pi starvation, seeds of soybean genotype HN89 were geminated in sand prior to selecting uniform seedlings after 5 d. Selected seedlings were then inoculated with highly effective rhizobium strain BXYD3 by immersing roots in a rhizobial suspension for 1.5 h and transplanted into a -N (530 μM N) nutrient solution and treated with 5 μM (-P) or 250 μM (+P) KH_2_PO_4_, respectively (Li et al., 2015). Nodules, roots, and entirely expanded young leaves were harvested separately 7 and 14 days after inoculation (DAI). For analysis of *GmPTF1* expression in roots responding to P deficiency, uniform seedlings were transplanted into hydroponic systems treated with -P and +P nutrient solutions as described above for 25 days. All samples were stored at −80°C prior to RNA extraction and RT-qPCR analysis.

### RNA Extraction and RT-qPCR Analysis

Total high-quality RNA was extracted from soybean nodules, roots, and leaves using RNAisoTM Plus reagent (Takara Bio, Otsu, Shiga, Japan) according to the manufacturer’s instructions. Subsequently, all RNA samples were treated with RNase-free *DNase* I (TaKaRa, Japan) to remove genomic DNA. About 1 μg of RNA was used for first-strand cDNA synthesis using oligo d (T), dNTPs, and MMLV-reverse transcriptase (Promega, Madison WI, USA) based on the protocol from the supplier. RT-qPCR was performed using a LightCycler96 (Roche Diagnostics GmbH, Germany) with the 20 μL reaction volume containing 2 μL of 1:50 diluted cDNA, 0.6 μL of specific primers, 6.8 μL of ddH_2_O, and 10 μL of Trans Start Top Green qPCR SuperMix (Trans). The reaction conditions for thermal cycling were as follows: 95°C for 1 min, 40 cycles of 95°C for 15 s, 60°C for 15 s, and 72°C for 30 s. Fluorescence data were collected during the step at 72°C. The housekeeping gene *EF-1α* from soybean (*TefS1*, accession no. X56856) or from tobacco (*Nicotiana tabacum; NtEF1a*, accession no. AF120093) (Schmidt and Delaney, 2010) was used as a reference gene to evaluate relative expression values. Relative expression was calculated as the ratio of the expression value of the target gene to that of *TefS1* or *NtEF1a* using the 2^-ΔΔCT^ method. All of the specific primers used for RT-qPCR are listed in Supplementary Table S1.

### Vector Construction

To characterize functional components of the *GmEXPB2* promoter, a series of deletions upstream of the translational start codon ATG were amplified by PCR. The six deletion fragments were named P1 (−304 to −1), P2 (−465 to −1), P3 (−687 to −1), P4 (−907 to −1), P5 (−1401 to −1), and P6 (−1799 to −1). These fragments were amplified using the common reverse primers P1-6-R and the forward primers P1-F, P2-F, P3-F, P4-F, P5-F, and P6-F. After digestion with *Eco*RI and *Bam*HI, the generated fragments were separately fused with a *GUS* reporter gene into the plant transformation vector pTF102.

To investigate the impact of E-box *cis*-elements located in the P1 region on *GUS* reporter gene expression, further deletion fragments were generated as pro1 (−213 to −1) and pro2 (−220 to −1) carrying one and two E-boxes, respectively. These fragments were amplified by PCR with the reverse primer P1-6-R and the forward primers pro1-F, and pro2-F. After verification by DNA sequencing, the pro1 and pro2 fragments were separately cloned into the pTF102 vector as described above.

For construction of plasmids with mutated E-box sequences, overlapping PCR was carried out first with the primers pro I-F/pro II-F/pro III-F and P1-6-R, as well as P1-F and pro I-R/pro II-R/pro III-R. Then, isolated sequence fragments were separately mixed and further used as templates to generate pro I, pro II, and pro III fragments with one, two or three mutated E-boxes amplified between the primers P1-F and P1-6-R. Among mutated fragments, the E-box sequence CATGTG in pro I was modified to ACTGGT, while the sequences CAATTG and GATGTG in pro II were respectively modified to AAATCG and ACTGTT, and the sequence CATTTG in pro III was modified to ACAAGT (highlight in blue, Supplementary Table S2).

To generate soybean transgenic composite plants overexpressing or suppressing *GmPTF1*, the open reading frame of *GmPTF1* was amplified using the *GmPTF1*-OE-F and *GmPTF1*-OE-R primers. After digestion with SwaI and BamHI, the fragment was cloned into the binary vector pFGC5941 with a 35S promoter. For the RNA interference construct, 337 bp of the *GmPTF1* coding sequence was amplified using the sense orientation primers *GmPTF1*-Ri-F1 and *GmPTF1*-Ri-R1 and the antisense orientation primers *GmPTF1*-Ri-F2 and *GmPTF1*-Ri-R2. The PCR products were digested separately and ligated into the SwaI and BamHI sites of the pFGC5941vector in the sense and antisense orientations. All primers used for the vector constructs are listed in Supplementary Table S2, and the restriction enzyme cutting sites are underlined in the corresponding primer sequences.

### Plant Transformation and Growth Conditions

The hypocotyl injection method was used to generate soybean transgenic composite plants as described previously (Kereszt et al., 2007), with some modifications. Hypocotyls of five-day-old seedlings with unfolded cotyledons were infected with *Agrobacterium rhizogenes* strain K599 carrying the target gene construct. Infected plants were grown in hydroponics under high humidity conditions. The main root was removed after hairy roots emerging from hypocotyl were approximately 10 cm long. Each individual hairy root was checked for green fluorescent protein (GFP) fluorescence to ensure the presence of the vector carrying target sequences. A single transgenic hairy root was kept for further study of each construct.

For histochemical analysis of *GUS* expression driven by the *GmEXPB2* promoter in hairy roots and nodules, transgenic soybean composite plants with hairy roots harboring *proGmEXPB2*::GUS constructs (Guo et al., 2011) were inoculated with rhizobia for 1 h and then transplanted into sand culture irrigated with -N (530 μM N) and -P (10 μM KH_2_PO_4_) solution for 7 and 14 days (Li *et al*., 2015).

For *GmEXPB2* promoter deletion analysis, transgenic soybean composite plants with hairy roots and harboring various truncated fragment constructs (P1-P6::GUS) were treated with high P (+P: 200 μM P added as KH_2_PO_4_) or low P (-P: 5 μM P added as KH_2_PO_4_) nutrient solutions for 15 days. All transgenic composite soybean plants were grown in a growth chamber with a 16 h/8 h, 26°C/24°C, light/dark photoperiod.

For transformation of tobacco leaves, recombinant plasmids were introduced into *Agrobacterium rhizogenes* strain EHA105 and then transiently transferred into tobacco leaves by infiltration. After 2 d, transgenic leaves were harvested for GUS staining, RNA extraction and fluorometric GUS assays.

### Histochemical GUS Staining and Fluorometric GUS Activity Assay

For histochemical analysis of *GUS* expression, all samples including soybean transgenic hairy roots and tobacco leaves were incubated in GUS staining solution containing 50 mM inorganic phosphate-buffered saline (Na_2_HPO_4_-NaH_2_PO_4_ buffer, pH 7.2), 0.1% (v/v) Triton X-100, 2 mM K_3_Fe(CN)_6_, 2 mM K_4_[Fe(CN)_6_]·3H_2_O, 10 mM EDTA-2Na, and 2 mM 5-bromo-4-chloro-3-indolyl-*β*-d-GlcA at 37°C for 24 h. After washing three times with 75% ethanol, stained samples were observed with a light microscope (Axio Imager A2m; Zeiss).

For the fluorometric GUS assay, transgenic soybean hairy roots and tobacco leaves were used to determine GUS enzyme activity by measuring the fluorescence of 4-methylumbelliferone (4-MU) produced by GUS cleavage of 4-methylumbelliferyl-*β*-d-glucuronide (4-MUG, Sigma, USA) according to the published procedure as described previously (Jefferson, 1988; Jefferson et al., 1987). Protein was extracted and quantified based on a published methods using bovine serum albumin as a standard (Bradford, 1976). Fluorescence was measured with a fluorescence spectrophotometer (HITACHI F-4600, Japan) at the excitation and emission wavelengths of 365 nm and 455 nm, respectively. GUS activity was calculated as nmol of 4-MU per minute per mg of protein.

### Database Search and Molecular Sequence Analysis

For analysis of *cis*-elements in the *GmEXPB2* promoter region, a 304 bp segment upstream of the translational start codon ATG was searched to locate potential *cis*-acting elements using NEW PLACE (https://www.dna.affrc.go.jp/PLACE/?action=newplace). The putative *cis*-elements are listed in Supplementary Table S3. For phylogenetic tree construction of GmPTFs, the basic helix-loop-helix (bHLH) transcription factors in soybean, a BLAST search was conducted at the phytozome website (http://www.phytozome.net), which yielded 170 bHLH genes in the soybean genome. Then, the phylogenetic tree was constructed based on whole protein sequence alignments using ClustalX and the neighbor-joining method with 1,000 bootstrap replicates in the MEGA 4.1 program.

### Subcellular Localization of GmPTF1 in Tobacco Cells

Subcellular localization of GmPTF1 was determined via transient expression of translational fusions with GFP in tobacco leave. First, the coding region of GmPTF1 was amplified using the specific primers GmPTF1-GFP-F and GmPTF1-GFP-R as listed in Supplementary Table S2. The resulting fragment of *GmPTF1* was then inserted into a modified pFGC5941 vector along with a *GFP* reporter gene after digestion by AscI. After transformation, the *Agrobacterium tumefaciens* strain EHA105 harboring 35S::GmPTF1-GFP or the 35S::GFP control vector was cultured in Luria-Bertani medium overnight. After centrifugation, the bacteria were re-suspended in infiltration medium (10 mM MgCl_2_, 10 mM MES, and 150 mM acetosyringone) to an OD600 of 0.45-0.55. Then, this suspension of cells containing GmPTF1-GFP constructs and the 35S empty vector was used to infiltrate leaves of 3-week-old tobacco plants. Infiltrated tobacco plants were grown for another 2 d and GFP florescence was observed using a confocal scanning microscope (LSM880; Carl Zeiss) with 488 nm excitation and 500-to 525-nm emission filter wavelengths.

### Measurement of Plant P Content

Soybean transgenic composite plants were dried at 105°C for 30 min, and then oven-dried at 75°C prior to weighting. About 0.2 g of dried sample was digested and total P content was measured using a continuous flow analyzer (SAN++). The resulting signals were analyzed in FlowAccess software (SAN++ FlowAccess V3 data acquisition Windows software package).

### Date Analysis

Results from RT-qPCR were normalized in each experiment. All data were statistically analyzed using Sigma Plot to calculate means and SE. Tests for statistical significance between groups were performed using Student’s *t* tests or a two-way ANOVA test in SPSS (version 17.0).

## Supplemental Data

**Supplemental Figure S1.** Effects of *GmEXPB2* on soybean pod number (A) and 100-grain weight (B) in field trials.

**Supplemental Figure S2.** Molecular identification of *GmEXPB2* in stable soybean transgenic lines.

**Supplemental Figure S3.** Phylogenetic analysis of GmPTF family members in soybean.

**Supplemental Figure S4.** Confirmation of RNA interference (Ri) and over-expression (OE) of *GmPTF1* in soybean transgenic composite plants.

**Supplemental Figure S5.** Effects of RNA interference (Ri) and over-expression (OE) of *GmPTF1* on root growth of soybean transgenic composite plants.

**Supplemental Table S1.** Gene-specific primers used for qRT-PCR analysis.

**Supplemental Table S2.** Gene specific primers used for *GmEXPB2* vector constructs.

**Supplemental Table S3.** List of *cis*-elements in the *GmEXPB2* promoter between the translation start codon and 304 bp upstream of the start codon.

## Acknowledgements

We thank Dr. Xiurong Wang for providing the seeds of overexpressing *GmEXPB2* soybean lines, and Thomas Walk of Golden Fidelity LLC for critical reading. The authors have no conflict of interest to declare.

## Notes

**Funding Information:** This work was financially supported by grants from the China National Key Program for Research and Development (2016YFD0100700), the National Natural Science Foundation of China (31601814), and the China Postdoctoral Science Foundation (2016T90592).

